# Extracellular matrix stiffness regulates fibroblast differentiation by influencing DNA methyltransferase 1 expression through microtubule polymerization

**DOI:** 10.1101/2022.10.20.513009

**Authors:** Zhihan Zhao, Guotao Huang, Yong He, Xiaohu Zuo, Wuyue Han, Hong Li

## Abstract

Cells sense physical cues, such as changes in extracellular matrix (ECM) stiffness, and translate these stimuli into biochemical signals that control various aspects of cellular behavior, thereby facilitating physiological and pathological processes in various organs. Evidence from multiple studies suggests that the anterior vaginal wall stiffness is higher in POP patients than in non-POP patients. Our experiments found that the expression of α-smooth muscle actin (α-SMA) in the anterior vaginal wall of patients with POP was increased, and the expression of DNMT1 was decreased. We used polyacrylamide gel to simulate matrix stiffening in vitro, and substrate stiffening induced the high expression of myofibroblast markers α-SMA and CTGF in L929 cells. Inhibition of DNMT1 promotes fibroblast differentiation into myofibroblasts in vitro. The results of bioinformatics analysis showed that the expression of DNMT1 was significantly correlated with microtubule polymerization-related proteins. The experiment showed that the microtubule polymerization inhibitor nocodazole could eliminate the decrease of DNMT1 expression in fibroblasts induced by high stiffness. We conclude that fibroblasts sense an increase in the stiffness of the surrounding matrix and regulate fibroblast differentiation by regulating the expression of DNA methyltransferase 1 (DNMT1) through the regulation of microtubule polymerization. This study may help to elucidate the complex crosstalk between vaginal fibroblasts and their surrounding matrix in both healthy and pathological conditions, and provide new insights into the implications of potentially targeted phenotypic regulation mechanisms in material-related therapeutic applications.

## 1. Introduction

Pelvic organ prolapse (POP) is an anatomical abnormality and pelvic organ dysfunction caused by pelvic floor support dysfunction[1, 2]. Epidemiological data showed that the prevalence of symptomatic POP among Chinese women was 9.1%[3, 4]. POP patients seriously affect the quality of life of patients. With the occurrence of global aging, the prevention and treatment of this disease is quiet indispensable.

Up to now, the pathogenesis of pelvic organ prolapse is still unclear, and one of the recognized major risk factors is stress injury[5], which involves extracellular matrix remodeling[6]. The extracellular matrix (ECM) is a three-dimensional network of acellular macromolecules composed of collagen, elastin, fibronectin, laminin, proteoglycans/glycosaminoglycans, etc[7]. Various components bind to each other through cell adhesion receptors to form a complex network. The ECM regulates multiple cellular functions, such as survival, growth, migration, and differentiation, by transducing signals through cell surface receptors into the cell surface, and is essential for maintaining normal homeostasis[8]. The ECM is a highly dynamic structural network that continuously undergoes remodeling mediated by several matrix-degrading enzymes under normal and pathological conditions, in addition to being regulated by fibroblasts, [9], which are widely present in connective tissue and are the main cells that regulate the components of extracellular matrix[10]. Differentiation of fibroblasts into myofibroblasts is a key feature of wound healing in soft tissues such as the vagina. When fibroblasts are grown in an environment with higher stiffness, some differentiate into myofibroblasts[11–13], which are marked by alpha smooth muscle actin (α-SMA)[14], connective tissue growth factor (CTGF) and type I collagen production. Myofibroblasts secrete collagen more vigorously and have contractile function, which can accelerate tissue repair and promote wound healing in the acute phase of tissue damage repair, but their long-term existence will cause excessive collagen deposition, tissue contracture, loss of elasticity, and dysfunction[15]. Mounting evidence suggests that the microenvironment of vaginal tissue in POP patients is more rigid than healthy vaginal tissue[14, 16], and POP mesh was also prone to a series of complications due to excessive stiffness[17, 18]. Therefore, it is of great clinical significance to study the effect of extracellular matrix stiffness on fibroblast fate regulation.

DNA methylation plays an important role in maintaining normal development and biological function, and covalent methylation of DNA cytosines promotes transcriptional repression in most cases[19]. The importance of DNA methylation is reflected in the development of many diseases due to DNA hypermethylation or hypomethylation and inappropriate temporal or spatial regulation. Maintenance of methylation patterns during cellular replication is mediated by DNA methyltransferase 1 (DNMT1), which catalyzes the transfer of methyl groups from S-adenosylmethionine to hemimethylated DNA[20, 21].

In this study, we found that the expression of α-SMA in the anterior vaginal wall of POP patients was significantly lower than that of control patients. Because the stiffness of the anterior vaginal wall in POP patients was higher than that in the control group, we cultured L929 cells with gels with different stiffnesses to simulate different extracellular matrix stiffnesses.

The morphology of fibroblasts in the gel was altered as the extracellular matrix stiffness increased.After 24 hours of culture in low stiffness gels, cells were more spherical, while after 24 hours of culture in high stiffness gels, cells were more fusiform and cells cultured in high stiffness gels had higher cell aspect ratios. The expression of DNMT1 in fibroblasts cultured in high-stiffness gel cells decreased, while the expression of fibroblast differentiation markers α-SMA and CTGF increased, indicating that fibroblasts cultured in high-stiffness gel cells differentiated into myofibroblasts. Subsequently, we performed immunohistochemical staining on the anterior vaginal wall of POP and control groups, and the results showed that the expression of DNMT1 in the anterior vaginal wall of POP patients was lower than that of the control group. To explore whether DNMT1 is involved in myofibroblast differentiation, we treated fibroblasts cultured with low stiffness conditions with DNMT1 inhibitor 5’-Aza, and found that the expression of DNMT1 was decreased and the expression of α-SMA and CTGF was increased. In order to explore how extracellular physical signals are transformed into intracellular biological signals, we performed biosignal analysis on DNMT1, and found that the expression of DNMT1 was significantly correlated with the expression of microtubule-related proteins. Therefore, we treated cells cultured in high-stiffness gels with the microtubule inhibitor nocodazole and found that the cells had increased DNMT1 expression and decreased α-SMA, CTGF expression. It indicates that the physical signal of stiffness is transformed into biological signal through cellular microtubules and affects the expression of DNMT1 in fibroblasts. However, the expression of α-SMA and CTGF in fibroblasts decreased after the cells cultured in the high-stiffness gel were treated with estrogen, indicating that estrogen could partially reverse the high-stiffness-induced myofibroblast differentiation. We show that extracellular matrix stiffness affects DNMT1 expression through microtubule aggregation and thus affects fibroblast-to-myofibroblast differentiation, and estrogen can partially reverse the effect of high-stiffness matrix on fibroblast differentiation. Our study sheds new light on the pathogenesis of POP and the selection of POP implant materials.

## 2. Materials and Methods

### 2.1 Cell Culture and Treatment

All cell experiments used L929 cells, cultured in RPMI Medium 1640 basic (1X) medium. In order to inhibit the expression of DNMT1, the cells were first cultured in the medium for 24h and then in medium containing 5-Aza for 24h.To inhibit myosin ATPase and microtubule assembly, after cultured on the gel for 6h, cells were incubated with nocodazole in medium for 5 h, followed by a medium change for 24 h. In order to observe the effect of estrogen on the differentiation of fibroblasts, the cells were cultured in the gel for 24 h, and then replaced with estrogen-containing medium for 24 h.

### 2.2 Human Specimen

In order to explore the differences in the expression of myofibroblasts in the anterior vaginal wall of POP patients and normal women, with the approval of the Ethics Committee(WDRY2021-K001) and the informed consent of the patients, the patients with III-degree POP who underwent hysterectomy and pelvic floor reconstruction were selected as the experimental group. Patients with other benign diseases who underwent total hysterectomy were the control group, and the anterior vaginal wall

### 2.3 Western blot

Cells were harvested using a spatula and washed twice with cold phosphate buffered saline (PBS). Then the cells were lysed in lysis (RIPA Buffer, Cocktail, phosphorylase inhibitor, PMSF), after collecting the protein, and the protein concentration was detected by BCA method for subsequent experiments. Antibodies used include rabbit polyclonal antibody (pAb) against DNMT1 (purchased from Cell Signaling Technology), rabbit polyclonal antibody against α-SMA, mouse polyclonal antibody against GADPH, rabbit polyclonal antibody against Collagen1, and rabbit polyclonal antibody against Collagen3 antibody (purchased from Proteintech). Estradiol was purchased from Sigma. Nocodazole and 5’-Aza were purchased from MedChemExpress (MCE).

### 2.4 Immunohistochemistry and immunofluorescence staining

Paraffin-embedded vaginal tissue sections were dewaxed at 65°C and antigen repair was performed at 100°C in citrate buffer solution (0.01 mol/ L, pH 6.0) for 30 min. After blocking with 5% bovine serum albumin (BSA) for 1 h, sections were incubated with anti-α-SMA antibody or anti-DNMT1 overnight at 4°C. After incubation with biotin-labeled secondary antibodies or fluorescent secondary antibodies for 1 h at 37°C, staining was then added and all stained slides were viewed under an Olympus BX50 microscope

### 2.5 CCK8 assay

Cell viability was estimated by CCK-8 assay (Seven, China). Approximately 104 cells were seeded in 96-well plates with 100 μl of medium per well. After 24 h of culture, different doses of Decitabine or Nocodazole or Estrogen were added. Incubate each well with 10 μg of CCK-8 solution for 2 hours in the dark before measuring absorbance at 450 nm using a microplate reader.

### 2.6 Preparation of Polyacrylamide Gel (PA Gel) Substrates

The preparation method of polyacrylamide gel (PA glue) substrate refers to the research of Justin R et al.[22], mainly by changing the ratio of acrylamide and methylene bisacrylamide to adjust the stiffness of PA glue, the corresponding stiffness is 4kpa and 50kpa. After mixing the prepared liquid, add it into parallel glass with an interval of 1.0 mm, and let it stand for about 1 hour until the PA glue solidifies. Wash the PA gel 2-3 times with PBS, cover the surface of the PA gel with photocrosslinker (Sulfo-SANPA, abcam, 0.5 mg/ml in PBS), irradiate it with UV light for 1 h, and wash the PA gel with PBS 2-3 times to wash off excess photocrosslinker. Collagen I (15 ug/mL, Biosharp) was spread on the surface of the gel and placed in a 4°C incubator overnight. The next day, the collagen solution was poured off, washed 2-3 times with PBS, dried overnight, and the collagen-coated PA gel could be used to inoculate cells.

### 2.7 Microarray Data Archive

The expression profile of the GSE18224 array was retrieved from the GEO database. GSE18224 collected Affymetrix RAE 430 2.0 gene chip array to detect the left ventricular transcriptome of 4 mice combined with three factors of gender, genotype and surgical method. The expression profile of the database is based on the GPL1261 platform. Download series matrix files and data table title descriptions from the GEO database to screen for genes with significant associations with DNMT1.

## 3. Results

### 3.1 Differences in the expression of myofibroblast-related proteins between pop patients and controls

The cellular composition of the vaginal wall is complex, mainly composed of fibroblasts and smooth muscle cells (SMCs), which play important roles in the extracellular integrity and mechanical stretching of the anterior vaginal wall. Compared with fibroblasts from non-prolapsed sites, vaginal fibroblasts from prolapsed tissue showed altered functional characteristics and altered expression of certain genes in vitro. In order to explore the difference in the expression of myofibroblasts in the anterior vaginal wall of POP patients and normal women, with the approval of the Ethics Committee of Wuhan Unibversity Renmin Hospital and the informed consent of the patients, the patients with III degree POP who underwent hysterectomy and pelvic floor reconstruction were selected as the experimental group. Patients with other benign gynecological diseases who underwent total hysterectomy were selected as the control group. During the operation, the anterior vaginal wall tissue was collected and placed in culture medium or fixed with paraformaldehyde. The myofibroblast marker α-SMA was detected by Western Blot and immunohistochemistry.. The results showed that the expression of α-SMA in the anterior vaginal wall of POP patients was significantly higher than that of controls (fig1a-c).

**Figure 1.**
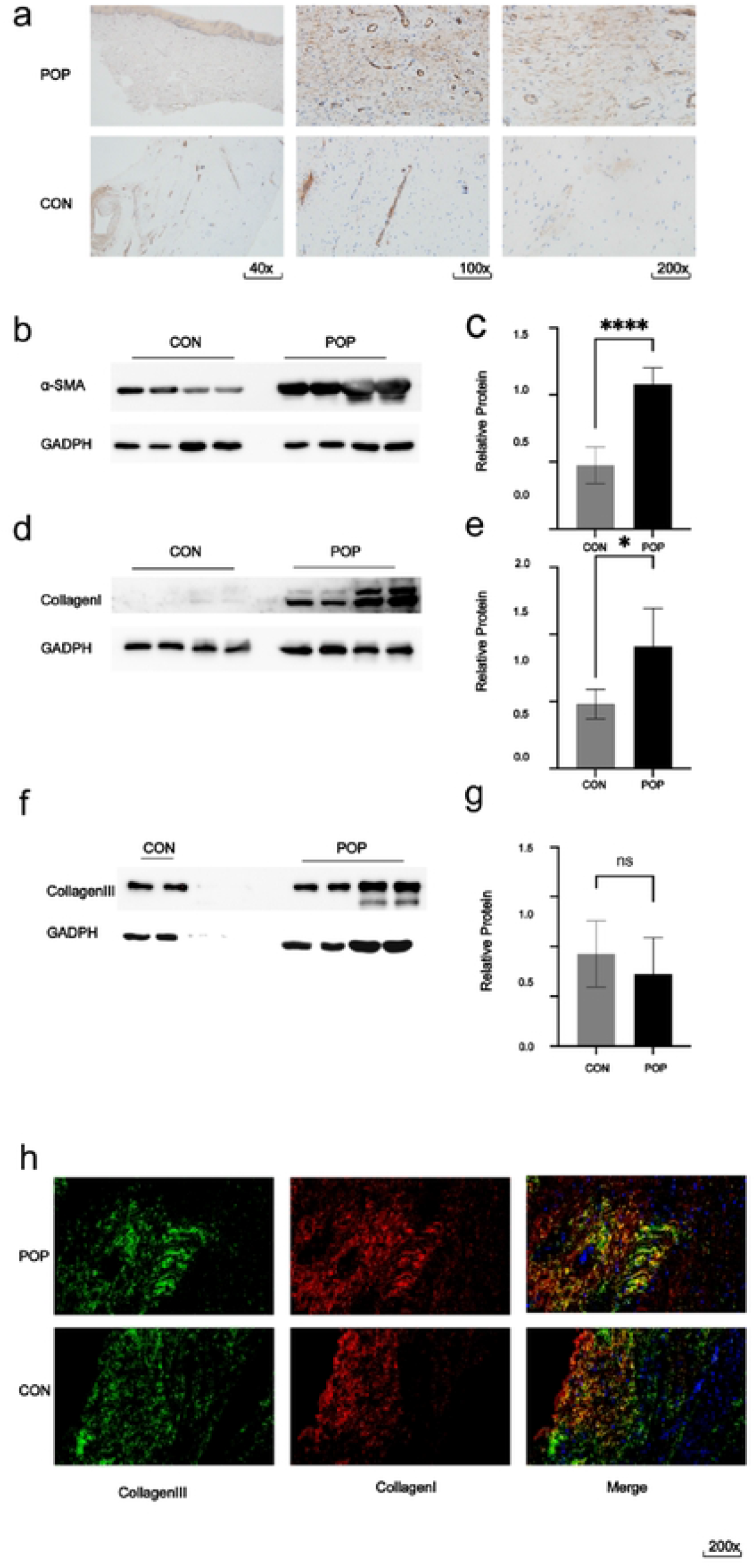
Differences in the expression of myofibroblast-related proteins between pop patients and controls (a) α-SMΛ immunohistochemical staining for control and POP patients; (b-c) Western blotting of α-SMA protein in control and POP patients.(d-g) Western blotting of Collagenl and CollagenIII protein in control and POP patients, (h) Immunofluorescence of CollagenI and CollagenIII protein in control and POP patients.

Myofibroblasts can secrete collagen, which can accelerate tissue repair and promote wound healing in the acute phase of tissue damage repair, but its long-term existence will cause excessive collagen deposition, tissue contracture, loss of elasticity, and dysfunction. Studies on changes in the number and proportions of collagen subtypes have yielded inconclusive data, both increases and decreases in total collagen content in the vaginal wall and pelvic floor supporting tissue have been reported in patients with POP. Collagen is the main component of the extracellular matrix of the anterior vaginal wall and is involved in the formation of the rigidity of the anterior vaginal wall, of which there are two main subtypes, type I and type III. We detected the expression of Collagen I and III in the POP group and the control group. The results of western blot and immunofluorescence showed that, compared with the control group, there was no significant difference in the expression of CollagenI, but the expression of CollagenIII increased in the POP group (fig1d-h).

### 3.2 Extracellular matrix stiffness affects fibroblast-to-myofibroblast transformation

The study found that the anterior vaginal wall tissue in women with POP was stiffer compared to healthy tissue, and the affected fibroblasts appeared to be affected by prolonged exposure to abnormally prolapsed ECM. During POP treatment, vaginal reconstruction using a polypropylene mesh that is stiffer than native tissue can further affect the prolapsed vaginal microenvironment. Stiffer meshes for the treatment of POP have been shown to produce more adverse events, and a recent animal model with POP showed that structural stiffness is a major driver of treatment failure. To investigate how extracellular matrix stiffness affects fibroblast fate determination and phenotypic regulation, we prepared PA gels with different stiffness (table1) and coated the surface of the photocrosslinker sulfo-sanpah, which was uniformly covered Collagen 1 (15ug/ml)after 1 h of activation under UV light. After 24 hours of adherent growth of L929 on the gel, the morphology was observed under an inverted microscope. We observed distinct morphological differences in cells grown in gels of different stiffnesses(fig2a). Cells grown on high-stiffness gels for 24 hours showed pleomorphic shapes such as star-shaped and long-spindle-shaped cells, while cells grown on low-stiffness gels were mainly round.Cell aspect ratio calculations indicated that cells cultured on high-stiffness gels had significantly higher aspect ratios than cells cultured on low-stiffness gels (Fig. b, c). Myofibroblasts are transient cells, their differentiation is reversible, they have contractile functions, and they secrete collagen more vigorously, so they can accelerate repair during acute tissue repair. Excessive deposition of collagen causes tissue contracture and loss of elasticity. Therefore, we cultured L929 under different stiffness for 48h and detected the expression of myofibroblast marker α-SMA. Western Blot results showed that cells cultured in high-stiffness gels had increased α-SMA expression (Fig. 2d,e).

**Fig.2.**
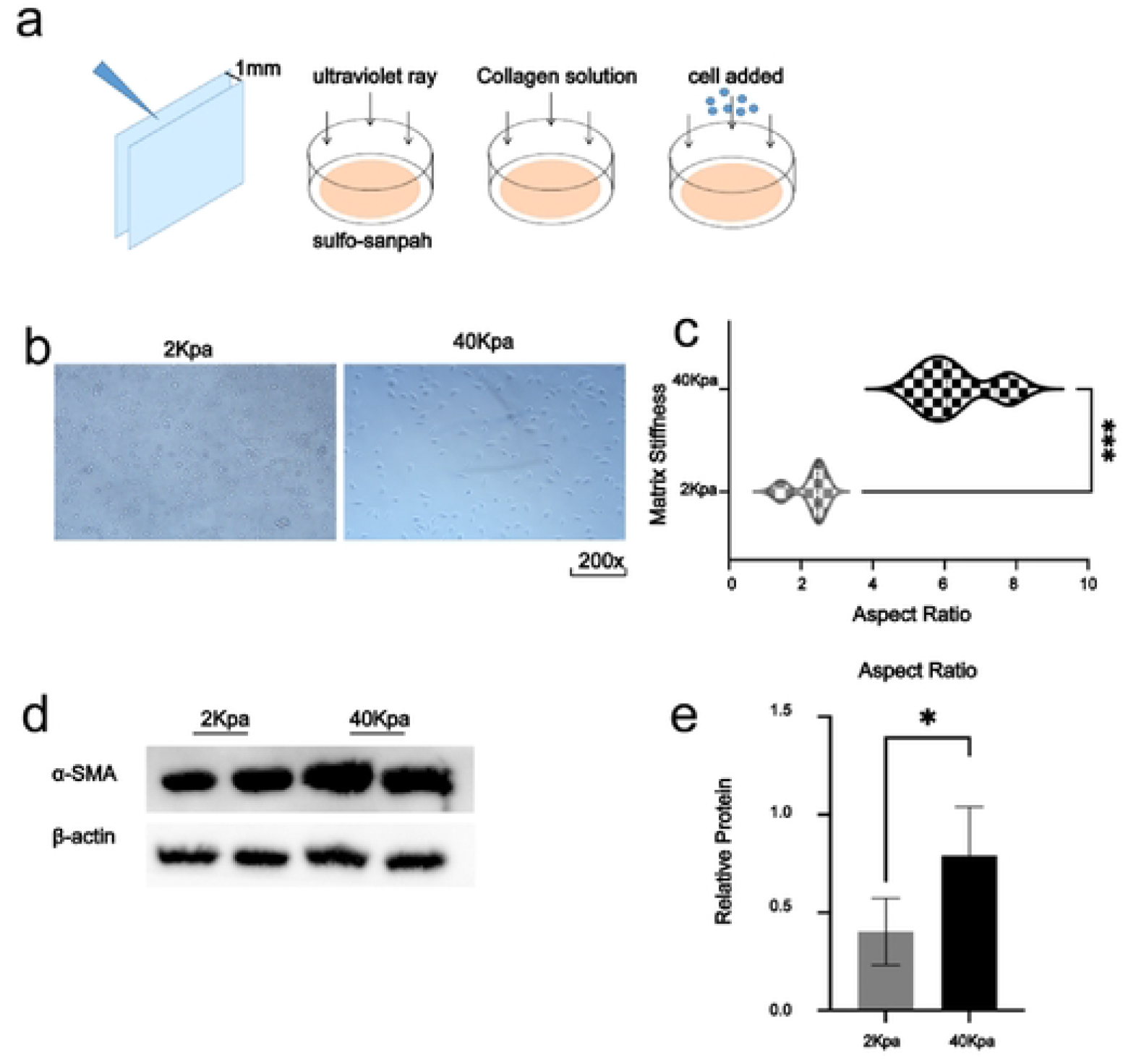
Extracellular matrix stiffness affects fibroblast-to-myofibroblast transformation (a)Flow chart of medium gel production, (b-c) Orthotopic microscope for cell culture; (d-e) Western blotting results of α-SMA protein

**Table 1.**

| Acrylamide % | Bisacrylamide% | Acrylamide<br>from 40% stock<br>solution (ml) | Bis-acrylamide<br>from 2% stock<br>solution (ml) | Water<br>(ml) | E ± St. Dev.<br>(kPa) |
| --- | --- | --- | --- | --- | --- |
| 4 | 0.3 | 1 | 1.5 | 7.5 | 2.24±0.58 |
| 8 | 0.48 | 2 | 2.4 | 5.6 | 40.4±2.39 |

### 3.3 Extracellular matrix stiffness regulates fibroblast-to-myofibroblast transition through DNMT1

While studies have demonstrated a critical role for epigenetic mechanisms (DNA methylation and demethylation, histone modifications, and RNA-based mechanisms) in mediating environmental signal-induced regulation of smooth muscle phenotypes Whether DNMT1 is involved in the transformation of fibroblasts to myofibroblasts and whether it is regulated by extracellular matrix stiffness. Therefore, we first detected the expression of DNMT1 in the anterior vaginal wall of POP patients and controls. Western blot results showed that the expression of DNMT1 in the anterior vaginal wall of POP patients was decreased (fig3a,b). Immunohistochemistry verified the above results (fig3c). Subsequently, we detected the expression of DNMT1 in fibroblasts grown in gels with different stiffnesses. After culturing L929 in gels with different stiffnesses for 48h, we collected the cells, extracted proteins, and performed western blot detection. The results showed that DNMT1 levels were significantly reduced after cells were cultured on high stiffness gels for 48 h (fig3d,f).To explore whether extracellular matrix stiffness affects fibroblast-to-myofibroblast differentiation through DNMT1, we added DNMT1 inhibitor to low-stiffness gel medium. We tested the chemosensitivity of L929 to Decitabine treatment by the CCK8 assay. The concentration gradient of decitabine is 0.5 ~ 10umol/L, and the final treatment concentration is 5umol/L. After the cells were cultured in the low-stiffness gel for 24 hours, the DNMT1 inhibitor Decitabine was added, and the medium was changed after 24 hours of culture, and the culture was continued for 24 hours. Western blotting results showed that compared with the control group, the expression of DNMT1 in the DNMT1 inhibitor group was significantly decreased, and the expression of α-SMA and CTGF was increased (fig3e, f).

**Fig3.**
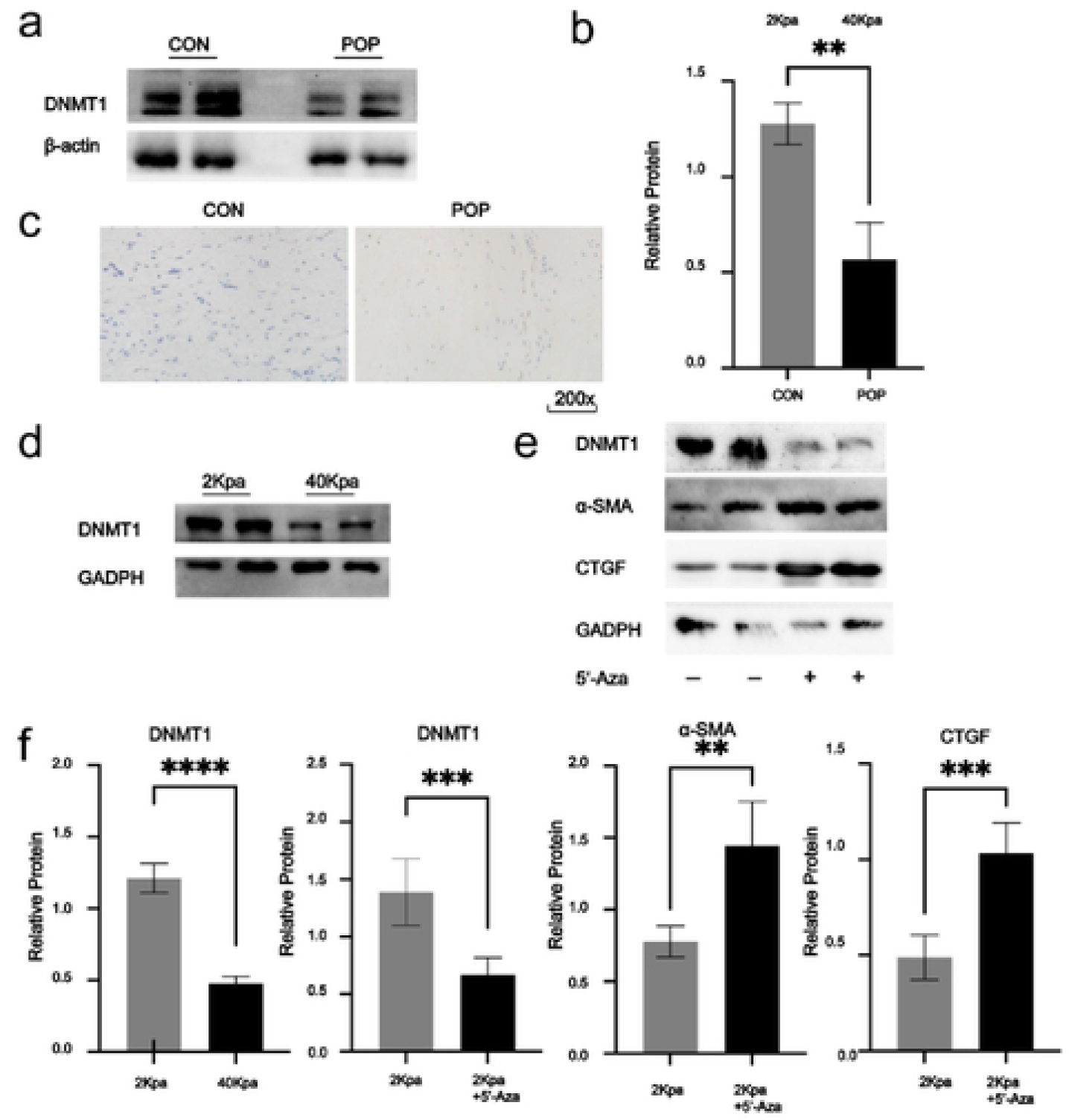
Extracellular matrix stiffness regulates fibroblast-to-myofibroblast transition through DNMT1 (a-c) Western blot and immunohistochemical staining of DNMT1 protein in the anterior vaginal wall of control and POP patients; (d,f) Expression of DNMT1 in cells cultured on gel with different stiffness.(e,f) The protein levels of these genes were detected by Western blot. Cells were treated with 5 ‘-AZA or PBS as control group and cultured on soft PA gel.

### 3.4 Cells transform physical signals into biological signals through the microtubule cytoskeleton

How extracellular matrix stiffness converts physical signals into intracellular biological signals is still unclear. It is known that cellular microtubules have kinetic properties of polymerization and depolymerization, and are involved in the maintenance of cell shape, cell division, signal transduction, and material transport. play an important role. Therefore, we retrieved the expression profile of the GSE18224 array from the GEO database. GSE18224 collected Affymetrix RAE 430 2.0 gene chip array to detect the left ventricular transcriptome of 4 mice combined with three factors of gender, genotype and surgical method. The expression profile of the database is based on the GPL1261 platform. The series matrix file and data table title descriptions of the database downloaded from the GEO database were used to screen genes with significant associations with DNMT1 using R software. The results showed that the microtubule-associated proteins Rmdn1 and Mast2 were significantly correlated with DNMT1, but not with the actin backbone (figure4a). We therefore treated cells with nocodazole, a rapidly reversible inhibitor of microtubule polymerization, and we tested the chemosensitivity of L929 to nocodazole treatment by CCK8 assay.The nocodazole concentration gradient is 1~100umol/L, and the final treatment concentration is 5umol/L. After culturing on the high stiffness gel for 6 hours, the cells are incubated in the medium containing nocodazole for 5 hours, and then renewed medium and the cells were cultured for an additional 24 hours. After 24 hours of adherent growth of L929 on the gel, the morphology was observed under an inverted microscope. We observed distinct morphological differences in cells grown in gels of different stiffnesses. Cells grown on high stiffness gels for 24 h showed pleomorphisms such as star-shaped and long-spindle, while the aspect ratio of cells decreased after the addition of nocodazole (Fig. 4b,c). Western blotting results showed that nocodazole treatment abolished the down-regulation of DNMT1 expression caused by high-stiffness gels and decreased α-SMA expression (fig4d,e), suggesting the role of the microtubule cytoskeleton in gel stiffness regulation of DNMT1 expression. potential effect.

**Figure 4.**
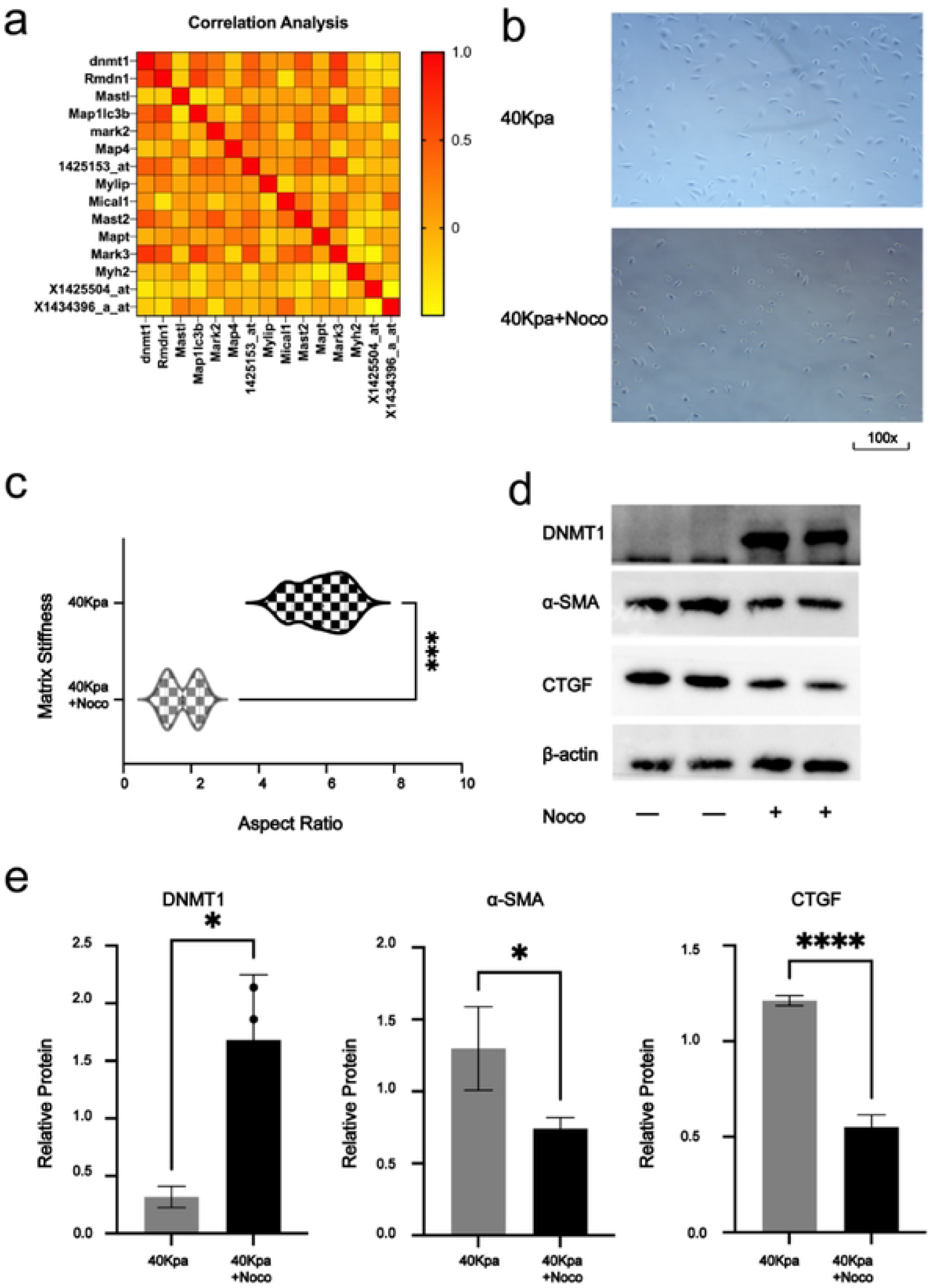
Cells convert physical signals to biological signals through the microtubule cytoskeleton(a) Gene correlation analysis showed that DNMT1 expression was significantly correlated with genes related to microtubule polymerization; (b,c) The morphological differences of cells cultured with high stiffness gel and added with Nocodazole were observed under forward microscope.(d,e) The protein levels of these genes were detected by Western blot. Cells were treated with 5 ‘-AZA or PBS as control group and cultured on soft PA gel.

### 3.5 Estrogen partially inhibits fibroblast differentiation induced by high stiffness gel

Among the clinical treatment options for POP, local vaginal estrogen administration after surgical treatment, studies have shown that there are significant gender differences in cardiac fibrosis. Therefore, we explored whether estrogen has an inhibitory effect on fibroblast differentiation. By adding estradiol to the medium, we tested the chemosensitivity of L929 to estradiol treatment by CCK8 assay. The estradiol concentration gradient was 10-8~10-5mol/L, and the final treatment concentration was 10-7mol/L. After the cells were cultured on gels of different hardness for 24 h, estradiol was added for 24 h, and the expressions of SMA and DNMT1 in L929 cells were detected. The results showed that cells cultured in high stiffness gels exhibited decreased differentiation and decreased cell aspect ratio after estrogen treatment compared to treatment without estrogen (Fig. 5a,b). Western blot results showed that the expression of α-SMA and CTGF in cells cultured on high-stiffness gels decreased after estrogen treatment (Fig. 5c,e), and the expression of DNMT1 increased, indicating an inhibition of fibroblast-to-myofibroblast differentiation effect (Fig. 5d,e).

**Figure 5.**
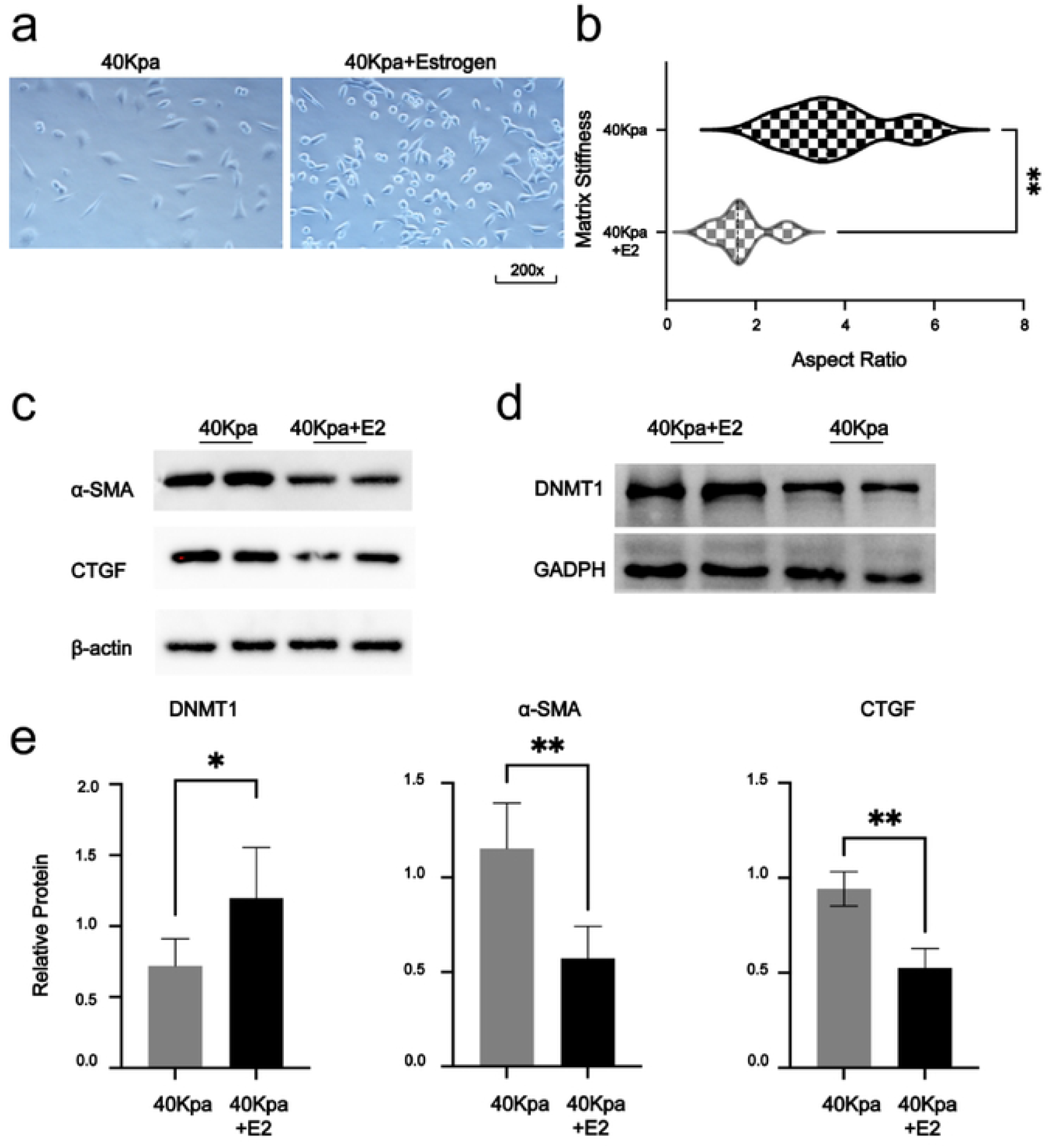
Estrogen partially inhibits fibroblast differentiation induced by high stiffness gel (a, b) The cell morphology was observed under a forward microscope, and the fiber cells composed of 4Okpa gel + estrogen were mostly round in shape, with a lower aspect ratio.;(c-e). Western blotting results of 4Okpa group and 4Okpa÷ estrogen group.

## 4. Discussion

Fibroblasts can sense an increase in environmental stiffness and fibroblast-to-myofibroblast differentiation occurs, a process known as fibroblast-myofibroblast transformation. Myofibroblasts have a stronger secretory function, which can accelerate tissue repair during tissue damage repair, but continuous activation can cause pathological deposition of extracellular matrix and cause tissue loss of elasticity. It is known that under physiological conditions, fibroblasts in the anterior vaginal wall of women are relatively quiescent and are characterized by low levels of α-SMA expression. However, when we detected the level of α-SMA in the anterior vaginal wall of POP patients and the control group, we found that the expression of α-SMA in the anterior vaginal wall of POP patients was significantly higher than that of the control group, indicating that myofibroblasts were continuously activated in the anterior vaginal wall of POP patients. How it came about is still unknown. Several studies have shown that the elastic modulus of the anterior vaginal wall in POP patients is significantly higher than that of the control group, that is, the extracellular matrix stiffness of the anterior vaginal wall is increased. In recent years, emerging evidence has emerged that the epigenetic mechanism of vascular smooth muscle cells (DNA methyl methylation and demethylation, histone modifications, and RNA-based mechanisms) play critical roles in mediating environmental stiffness signaling to induce various cellular phenotypic regulation. However, the understanding of epigenetic regulation triggered by mechanical stimulation of fibroblasts is limited. Here, we report the effect of microenvironmental stiffness on fibroblast differentiation mediated by DNMT1. DNMT1 is a mechanosensitive protein, and our study uncovers a fundamental role for DNMT1 as a mechanosensor downstream element in how fibroblasts sense their physical microenvironment. When the matrix becomes elastically stiff, cells produce less DNMT1, resulting in lower levels of DNA methylation. The detailed mechanotransduction mechanism by which ECM stiffness regulates DNMT1 awaits further characterization, but our experimental results suggest that the process involves the aggregation of microtubules. These findings have potential implications for the study of mechanotransduction and underscore the importance of DNMT1 in mediating the interaction between fibroblasts and matrix stiffness. In addition, in the routine clinical treatment of POP patients after surgery, there is local application of estrogen to the vagina, and studies have shown that estrogen combined with biofeedback electrical stimulation therapy improves the degree of prolapse compared with the biofeedback electrical stimulation group. Here we explored the possibility that estrogen affects fibroblast differentiation in stiffness and showed that estrogen treatment increased DNMT1 expression and decreased α-SMA expression in fibroblasts. However, this study still has some limitations. First, we have not yet determined how physical signals are converted into biological signals through cellular microtubule aggregation. Our preliminary study found that integrin family and Hippo family may be involved in this process. Second, how estrogen regulates the expression of DNMT1 remains to be explored. Preliminary studies suggest that estrogen may affect the expression of DNMT1 by interacting with ER2. Our study provides an explanation for the underlying mechanism of fibroblast differentiation by showing that when a mechanosensitive protein (DNMT1 in our case) receives signals from the mechanical microenvironment, cells respond through changes in their stiffness, and provides new ideas and possibilities for the treatment of POP patients and the selection of stiffness-related properties of implant materials.

